# Multi-environment analysis enhances genomic prediction accuracy of agronomic traits in sesame

**DOI:** 10.1101/2022.11.26.518043

**Authors:** Idan Sabag, Ye Bi, Zvi Peleg, Gota Morota

## Abstract

Sesame is an ancient oilseed crop containing many valuable nutritional components. Recently, the demand for sesame seeds and their products has increased worldwide, making it necessary to enhance the development of high-yielding cultivars. One approach to enhance genetic gain in breeding programs is genomic selection. However, studies on genomic selection and genomic prediction in sesame are limited. In this study, we performed genomic prediction for agronomic traits using the phenotypes and genotypes of a sesame diversity panel grown under Mediterranean climatic conditions over two growing seasons. We aimed to assess the accuracy of prediction for nine important agronomic traits in sesame using single- and multi-environment analyses. In single-environment analysis, genomic best linear unbiased prediction, BayesB, BayesC, and reproducing kernel Hilbert spaces models showed no substantial differences. The average prediction accuracy of the nine traits across these models ranged from 0.39–0.79 for both growing seasons. In the multi-environment analysis, the marker-by-environment interaction model, which decomposed the marker effects into components shared across environments and environment-specific deviations, improved the prediction accuracies for all traits by 15%–58% compared to the single-environment model, particularly when borrowing information from other environments was made possible. Our results showed that single-environment analysis produced moderate-to-high genomic prediction accuracy for agronomic traits in sesame. The multi-environment analysis further enhanced this accuracy by exploiting marker-by-environment interaction. We concluded that genomic prediction using multi-environmental trial data could improve efforts for breeding cultivars adapted to the semi-arid Mediterranean climate.

## Introduction

Sesame (*Sesamum indicum* L.) is an ancient oilseed crop with an annual global production of 6.8 million tons (https://www.fao.org/faostat/en/#data/QCL), and there is an increasing demand for its consumption because of its valuable nutritional components. Sesame seeds are rich in high-quality fatty acids, proteins, minerals, and antioxidants, which have health benefits (Wei et al., 2022). The recent availability of sesame genome resources (Berhe et al., 2021; Wang et al., 2022) has provided an opportunity for quantitative genetic modeling of sesame populations. For example, using these resources, quantitative trait loci mapping and genome-wide association analysis in sesame have been conducted for identifying its morphological traits (Mei et al., 2017; Sabag et al., 2021), yield components (Zhou et al., 2018; Sabag et al., 2021), plant architecture (Teboul et al., 2022), response to biotic (Asekova et al., 2021) and abiotic (Li et al., 2018; Dossa et al., 2019) stresses, and seed quality traits (Teboul et al., 2020; Cui et al., 2021) to understand the underlying genetic basis. However, little is known regarding the ability of genomics to predict genetic or breeding values in sesame. Complex traits are influenced by multiple genes, with small effects that are not statistically significant. To address this challenge, genomic predictions that simultaneously accommodate all available genetic markers in regression models to predict genetic or breeding values for capturing marker genetic effects across the whole-genome (Meuwissen et al., 2001) are being used. Genetic or breeding values of lines can be incorporated into selection indices to make a selection decision in breeding (Smith, 1936; Hazel, 1943).

Agronomic traits are influenced by genetic by environment interactions (G × E) (Gadri et al., 2020). The impact of G × E ranges from changes in the relative ranking of genotypes to the genomic prediction accuracy, making breeding decisions challenging. With the availability of whole-genome data, the factors of G × E can be reparametrized as functions of molecular genetic markers via marker-by-environment interactions (M × E). Recent efforts have included the use of M × E in whole-genome regression models (Lopez-Cruz et al., 2015; Crossa et al., 2016). These studies showed that modeling M × E could increase the prediction accuracy compared with that of models without the M × E term.

In this study, we used phenotypic and genomic data from a sesame diversity panel (SCHUJI panel) that was grown over two years (environments) under Mediterranean climatic conditions. This panel was recently used to perform genome-wide association analysis and estimate genomic heritability and genomic correlations for various agronomic traits (Sabag et al., 2021). Our study aimed to evaluate the utility of genomic prediction in predicting sesame traits for both single- and multi-environment analyses.

## Materials and Methods

### Plant materials, field experiments and genomic data

The complete dataset included phenotypic and genomic data of 182 sesame genotypes from the SCHUJI panel grown over two seasons (2018 and 2020) at the experimental farm of the Hebrew University of Jerusalem (Rehovot, Israel) (Sabag et al., 2021). This panel was characterized by nine agronomic traits: flowering date (FD, in days), height to the first capsule (HTFC, in cm), plant height (PH, in cm), reproductive zone (RZ, in cm), reproductive index (RI, a ratio), number of branches per plant (NBPP), seed-yield per plant (SYPP, g), seed number per plant (SNPP, in gm), and thousand-seed weight (TSW, in gm). The summary statistics for these traits are presented in Table S1. The best linear unbiased estimates of the genotypes were calculated per year by treating the block effect as random (Sabag et al., 2021). Genotyping by sequencing was used to obtain marker information for the 182 genotypes (Elshire et al., 2011). The quality control step included removing tightly linked markers (*r*^2^ ≥ 0.99), minor allele frequencies less than 0.05, and heterozygosity rates greater than 0.2. The remaining 20,294 single nucleotide polymorphism (SNPs) markers were used for subsequent analyses (Sabag et al., 2021).

## Statistical analyses

### Single-environment analysis

A single-environment analysis was conducted by fitting two kernel-based methods, genomic best linear unbiased prediction (GBLUP) (VanRaden, 2008) and reproducing kernel Hilbert spaces regression (RKHS) (de los Campos et al., 2010); and two variable selection methods, BayesB (Meuwissen et al., 2001) and BayesC (Kizilkaya et al., 2010).

The kernel-based methods GBLUP and RKHS were fitted as follows.

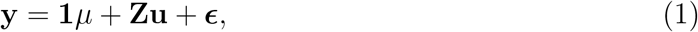

where **y** is the vector of phenotypes; **1** is the vector of ones; *μ* is the overall mean; **Z** is the incidence matrix for the random effect; 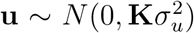 is the vector of random genotypes; and 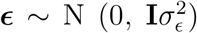 is the random residual effect. Here, the kernel matrix **K** was set to the genomic relationship matrix (**G**) and the Gaussian kernel matrix (**GK**) in GBLUP and RKHS, respectively; **I** is the identity matrix; 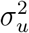 is the genetic variance; and 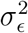 is the residual variance. The genomic relationship matrix captures additive gene action. In contrast, the Gaussian kernel is equivalent to a space continuous version of the diffusion kernel deployed on graphs (Morota et al., 2013), which can model additive by additive epistatic gene action up to an infinite order (Jiang and Reif, 2015). In GBLUP, 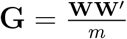, where **W** is a centered and standardized gene content matrix and *m* is the total number of SNP markers. The Gaussian kernel between a pair of lines *i* and *i*′ with their marker vectors **w**_*i*_ and 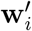 is given by

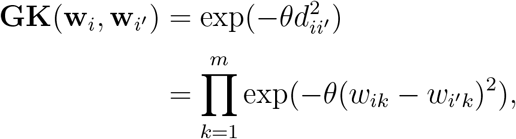

where 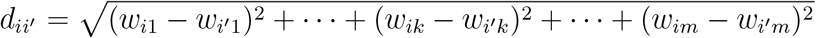 is the Euclidean distance and *θ* is the bandwidth parameter. Here, large *θ* leads to **GK** entries closer to 0 (i.e., local kernel), and smaller *θ* produces entries closer to 1 (i.e., global kernel), controlling the magnitude of genetic similarity between lines. The bandwidth parameter was determined using kernel averaging or multiple kernel learning (de los Campos et al., 2010) by fitting two contrasting kernel matrices with *θ* = 0.2 and 1.2.

The variable selection methods BayesC and BayesB followed

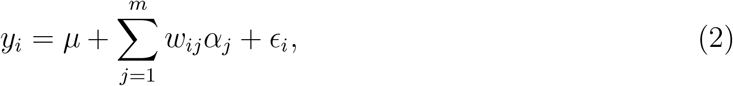

where *y*_*i*_ is the vector of phenotypes for the *i*th genotype; *μ* is the overall mean; *w*_*ij*_ is the marker covariate at the *j*th SNP marker coded as 0, 1, or 2; *m* is the number of SNPs; and *α*_*j*_ is the *j*the marker effect. The prior of *α*_*j*_ for BayesB was:

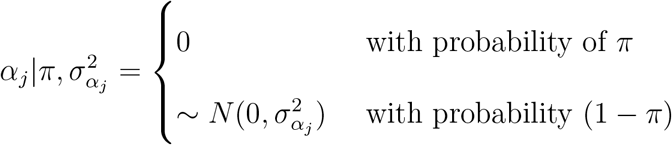

where 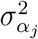 is the marker genetic variance for the *j*th SNP and *π* is a mixture proportion set to 0.99. A Gaussian prior 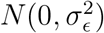 was assigned to the vector of residuals, and a flat prior was assigned to *μ*. The scaled inverse *χ*^2^ distribution was assigned to 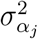 by setting the degrees of freedom equal to 5 and choosing the scale parameter, assuming that the model explained 50% of the phenotypic variance. In BayesC, 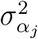 was replaced with the common marker genetic variance 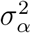.

#### Multi-environment analysis

A multi-environment analysis was conducted using the M × E model (Lopez-Cruz et al., 2015). The core idea of the M × E model is to partition the total marker genetic effects into the main marker genetic effects across all environments and specific marker effects in each environment. As a vector of genetic values consists of a linear combination of marker effects, G × E GBLUP is equivalent to M × E ridge regression BLUP (RR-BLUP). The M × E RR-BLUP model is expressed as 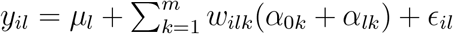, where 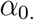 is the main effect of the markers stable for all the environments, 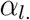 is the specific effect of the markers unique for each environment, and *l* is the *l*th environment. In matrix notation,

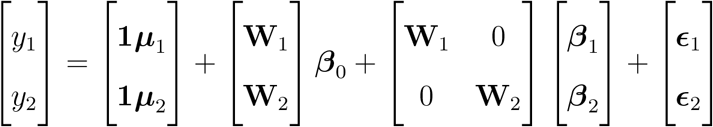

where 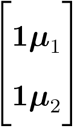 is the vector of grand means; 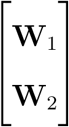 is the matrix of centered and standardized marker matrix for each environment; 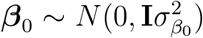 is the marker effects among environments; the variance component 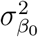 is common across the environments and borrows information among them; 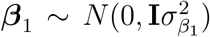 and 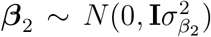 capture the environment specific marker effects with their environment specific variances; and 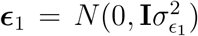 and 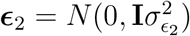 are the heterogeneous residual variances. The extent of variance components associated with ***β***_0_ relative to ***β***_1_ and ***β***_2_ suggests the importance of M × E. The grand mean was assigned a flat prior. The variance components of markers were drawn from a scaled inverse *χ*^2^ distribution with degrees of freedom *ν* = 5 and scale parameter *s* such that the prior means of variance components equal half of the phenotypic variance.

Additionally, the genomic correlation between the same trait in different environments was estimated using a bivariate GBLUP model by extending the single-environment variancecovariance structure to

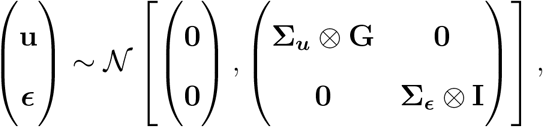

where **I** is an identity matrix and **Σ**_***u***_ and **Σ**_***ϵ***_ are genetic and residual variance-covariance matrices, respectively. Genomic correlations were derived as 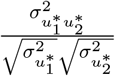 where 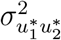 is the additive genetic covariance of the trait between the two environments, and 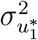 and 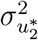 are additive genetic variances of the trait in 2018 and 2020, respectively. The covariance matrices, **Σ**_***u***_ and **Σ**_***ϵ***_, were assigned an inverse Wishart prior distribution with **W**^−1^(**S**_*u*_, *ν*_*u*_) and **W**^−1^(**S**_*ϵ*_, *ν*_*ϵ*_), respectively; **S**_*u*_ and **S**_*ϵ*_ are the identity matrices; and *ν*_*u*_ and *ν*_*ϵ*_ are the degrees of freedom. In addition, the phenotypic correlation between the two environments was estimated using the sample phenotypic correlation and the variance components obtained from the M × E model. The full data set was used to estimate the variance components and genetic correlations.

All the models were implemented in a Bayesian manner. Posterior inferences were based on 50,000 Markov chain Monte Carlo samples, 20,000 burn-in, and a thinning rate of 5 using the BGLR R package following default rules for choices of hyperparameters (Pérez and de Los Campos, 2014; Pérez-Rodríguez and de Los Campos, 2022).

### Cross-validation scenarios

For the single-environment analysis, the prediction accuracies of the GBLUP, BayesB, BayesC, and RKHS models were evaluated using the repeated random subsampling cross-validation (CV) (Figure 1). Two-thirds of the lines were used as a training set (TRN) and the remaining one-third were used as a testing set (TST). We measured the predictive Pearson correlation for each repeat, between the observed and predicted values in the TST. The average across 50 replications was used to derive the prediction accuracy of the model.

**Figure 1:**
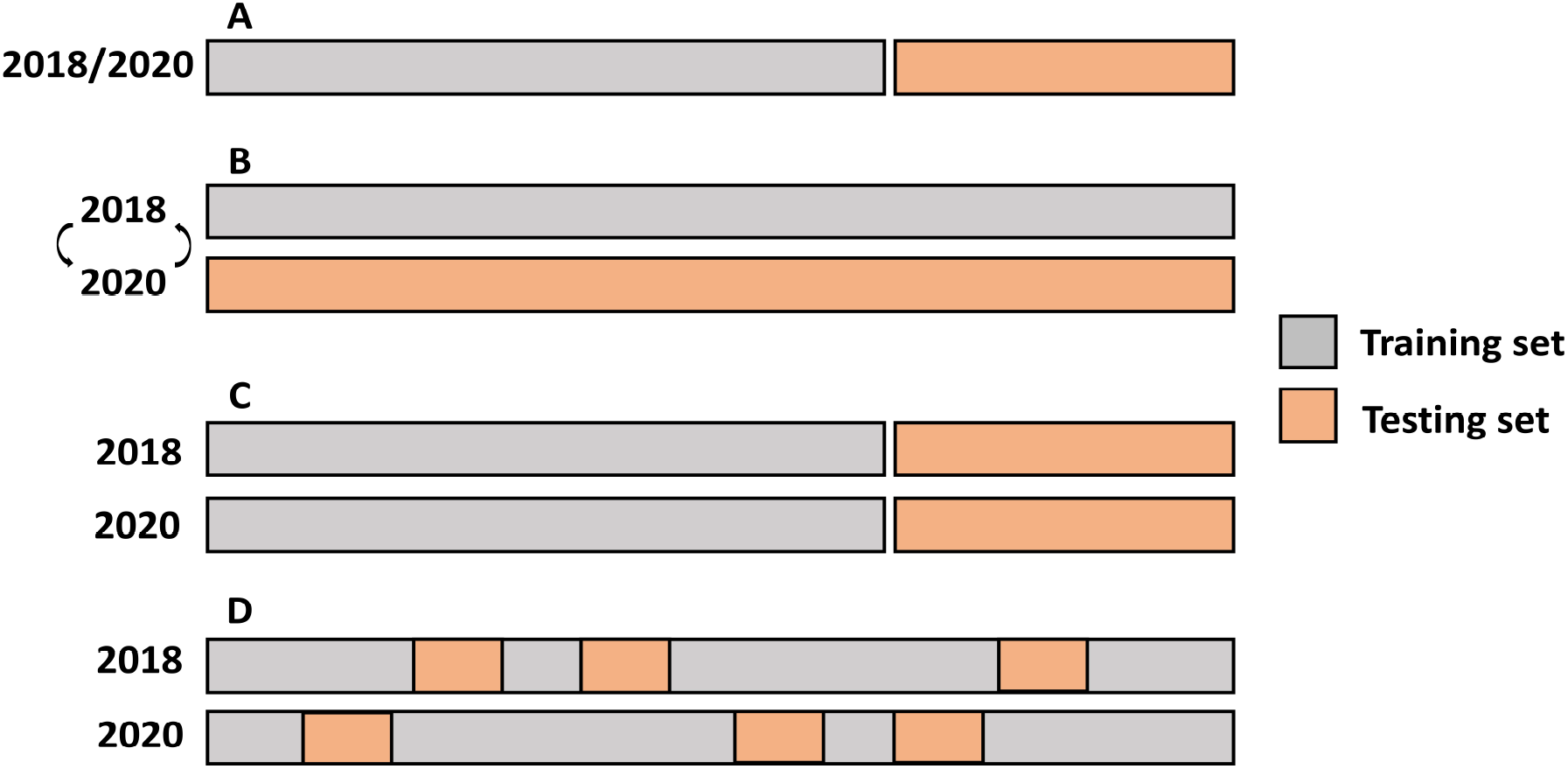
Single- and multi-environment genomic prediction cross-validation scenarios. A: Single-environment analysis, B: All the lines in one environment were used to predict the same lines in a new environment (CV0), C: Performance of new lines that are not phenotyped in any environment was predicted through the genetic relationship with other lines (CV1), and D: Predict lines that were evaluated in only one environment through the genetic and environmental relationships (CV2).

The predictive ability of the multi-year analysis was assessed using three different CV scenarios that simulated various prediction challenges in plant breeding (Burguenõ et al., 2012) (Figure 1). In the first scenario, leave one environment-out CV (CV0), used all the lines in one environment to predict the same lines in a new environment. The second scenario (CV1) predicted the performance of new lines that were not phenotyped in either environment. This scenario evaluated whether newly developed lines that had never been observed in any of the environments could be predicted from their genetic relationships with other lines. In this scenario, the same lines in the same environments were used as TRN, whereas the remaining lines were used for TST. The third CV scenario (CV2) posed the following challenge: some lines were evaluated in only one environment owing to the sparse field design. In this case, the prediction leveraged both genetic and environmental relationships. The GBLUP model was used to evaluate CV0, and the performance of the M × E RR-BLUP model was benchmarked with that of GBLUP in CV1 and CV2. The repeated random subsampling CV was employed for CV1 and CV2.

## Data availability

The phenotypic and genomic information can be found at https://figshare.com/s/94a222afca9423d0b1aa and https://figshare.com/s/a061d548a97237b51a61, respectively.

## Results

The sample phenotypic correlations between the environments were all positive, ranging from 0.50 (SNPP) to 0.96 (FD) (Table 1). Similarly, variance component-derived phenotypic correlations were all positive, ranging from 0.37 (SNPP) to 0.80 (FD) (Table 1). Genomic correlation estimates between the environments were all positive, ranging from 0.63 (SNPP) to 0.97 (FD) (Table 1).

**Table 1.** Genomic heritability estimates of the nine agronomic sesame traits (*h*^2^), genetic correlations (*r*_*g*_), sample phenotypic correlations (*r*_*y*_), and variance-components derived phenotypic correlations 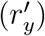 between the two environment using the marker-by-environment interaction model. Flowering date (FD), height to the first capsule (HTFC), plant height (PH), reproductive zone (RZ), reproductive index (RI), number of branches per plant (NBPP), seed-yield per plant (SYPP), seeds number per plant (SNPP), and thousand-seed weight (TSW).

| Trait | $h^2$ | $r_g$ | $r_y$ | $r'_y$ |
| --- | --- | --- | --- | --- |
| FD | 0.72 | 0.97 | 0.96 | 0.80 |
| HTFC | 0.68 | 0.94 | 0.95 | 0.77 |
| PH | 0.57 | 0.82 | 0.83 | 0.66 |
| RZ | 0.62 | 0.87 | 0.82 | 0.71 |
| RI | 0.68 | 0.92 | 0.93 | 0.75 |
| NBPP | 0.55 | 0.83 | 0.78 | 0.65 |
| SYPP | 0.38 | 0.76 | 0.58 | 0.47 |
| SNPP | 0.29 | 0.63 | 0.50 | 0.37 |
| TSW | 0.70 | 0.87 | 0.80 | 0.77 |

### Single-environment genomic prediction

Single-environment prediction accuracies of the nine agronomic traits were evaluated using the four whole-genome regression models (Figure 2 and Table S2). Overall, no notable difference was observed between the environments and the models. The highest mean prediction accuracy was obtained for HTFC (0.77 and 0.78 in 2018 and 2020, respectively, averaged across the models), whereas the lowest was for SNPP in 2018 (0.49) and SYPP in 2020 (0.39). FD, PH, RI, and NBPP showed relatively high prediction accuracies. In particular, the prediction accuracies ranged from 0.74 in 2018 to 0.70 in 2020 for FD, 0.68 in 2018 to 0.67 in 2020 for PH, 0.71 in 2018 to 0.74 in 2020 for RI, and 0.69 in 2018 to 0.62 in 2020 for NBPP. The prediction accuracy of RZ was slightly lower than that of these traits, with 0.56 in 2018 and 0.53 in 2020. The three yield-related traits SYPP, SNPP and TSW showed moderate prediction accuracies of 0.57 and 0.39, 0.49 and 0.40, and 0.55 and 0.50 for 2018 and 2020, respectively. The prediction accuracies for 2018 were higher than those for 2020.

**Figure 2:**
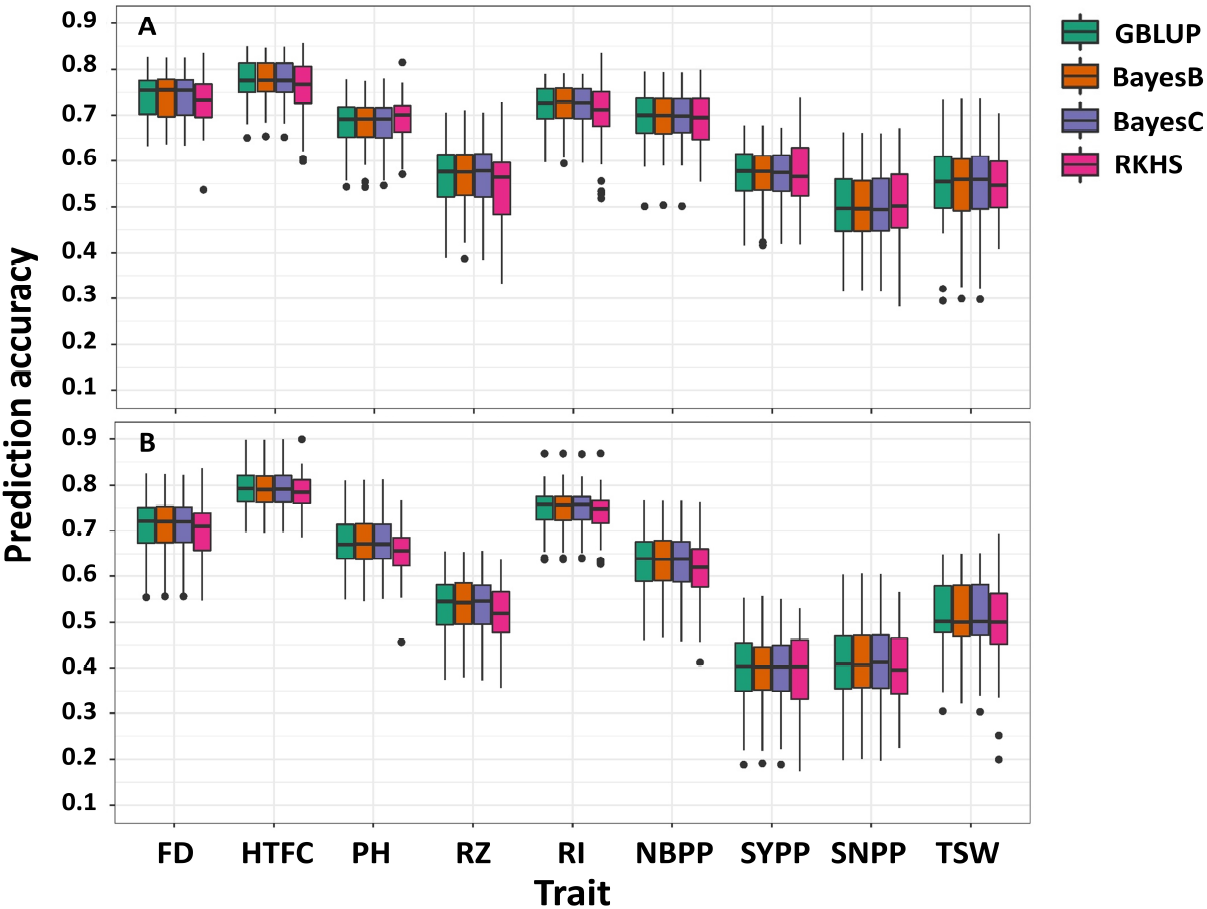
Single-environment prediction accuracies of the nine agronomic sesame traits in 2018 (**A**) and 2020 (**B**) growing seasons using genomic best linear unbiased prediction (GBLUP), BayesB, BayesC, and reproducing kernel Hilbert spaces regression (RKHS). Flowering date (FD), height to the first capsule (HTFC), plant height (PH), reproductive zone (RZ), reproductive index (RI), number of branches per plant (NBPP), seed-yield per plant (SYPP), seeds number per plant (SNPP), and thousand-seed weight (TSW).

### Multi-environment genetic parameter estimation

Variance component estimates were obtained from the M × E RR-BLUP model using the full data set and expressed in terms of proportions (Figure 3). In the two yield-related traits, SYPP and SNPP, the M × E components were largest whereas the additive genetic components were the lowest. However, the extent of M × E was lower for FD, HTFC, RI, and TSW. Similarly, the estimates of genomic heritability were low for SYPP and SNPP, and high for FD, HTPC, RI, and TSW (Table 1). Estimates of genomic correlations between the two environments were all moderate to high, ranging from 0.63 (SNPP) to 0.97 (FD) (Table 1).

**Figure 3:**
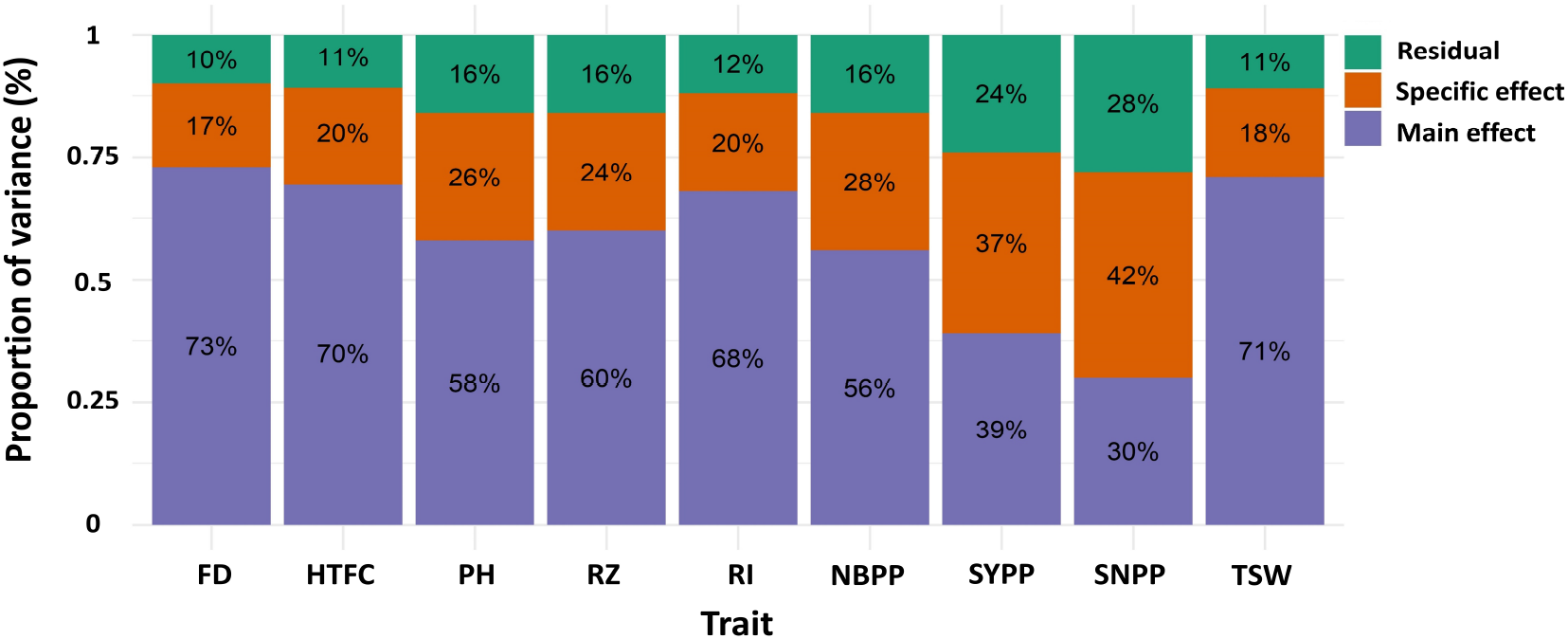
Proportion of the main genetic variance, environment-specific variance, and residual variance components for each trait obtained from the marker-by-environment interaction model. Flowering date (FD), height to the first capsule (HTFC), plant height (PH), reproductive zone (RZ), reproductive index (RI), number of branches per plant (NBPP), seed-yield per plant (SYPP), seeds number per plant (SNPP), and thousand-seed weight (TSW).

### Multi-environment genomic prediction

One of the main challenges for the genomic prediction of multi-environmental data was predicting the performance of new or observed lines in new or known environments. We used multi-environment data to evaluate the genomic prediction accuracies of nine important agronomic traits in sesame by accounting for M × E. Our main objective was to investigate whether obtaining information from another environment could improve predictions compared to a single-environment analysis. As we did not observe a difference among GBLUP, BayesB, BayesC, and RKHS in the single-environment analysis, multi-environment analysis was conducted using the GBLUP or RR-BLUP type of models.

#### CV0 scenario

In the CV0 scenario, all lines in one environment were used to predict the same lines in a new environment by applying the GBLUP model (Figure 1B). Overall, we obtained an improvement in the prediction accuracies of all traits compared to the single-environment model (Figure 4). The prediction accuracies were highest for FD and HTFC, with 0.93 and 0.92, respectively. For other agronomic traits, the prediction accuracies ranged between 0.78 (NBPP) and 0.9 (RI). For yield components, prediction accuracies were 0.63, 0.55, and 0.74 for SYPP, SNPP, and TSW, respectively.

**Figure 4:**
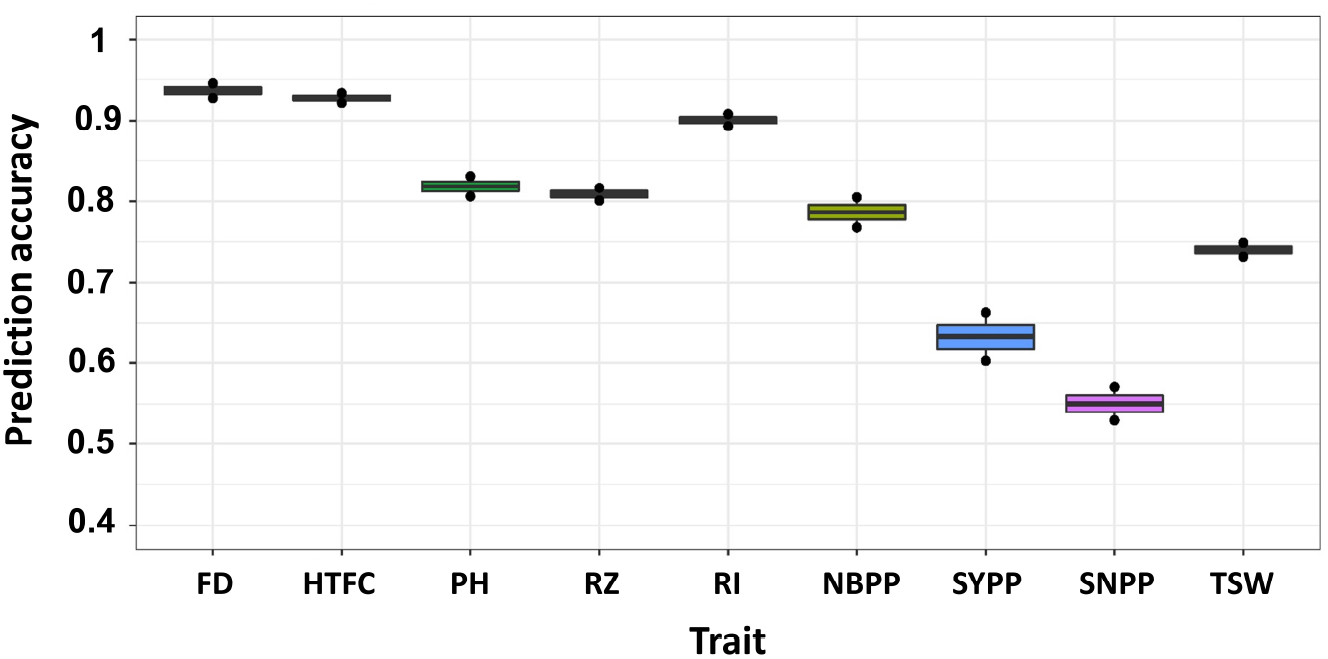
Multi-environment genomic prediction accuracies of the nine agronomic sesame traits using the best linear unbiased prediction model when all the lines in one environment were used to predict the same lines in a new environment (CV0). Flowering date (FD), height to the first capsule (HTFC), plant height (PH), reproductive zone (RZ), reproductive index (RI), number of branches per plant (NBPP), seed-yield per plant (SYPP), seeds number per plant (SNPP), and thousand-seed weight (TSW).

#### CV1 scenario

The CV1 scenario mimicked the situation in which we aimed to predict the performance of new lines (Figure 1C). We did not observe a major difference between the single-environment and M × E models (Figure 5 and Supplemental Table S3). The prediction accuracies from multi-environment analysis were almost equal to or lower than those from the single-environment analysis for some traits.

**Figure 5:**
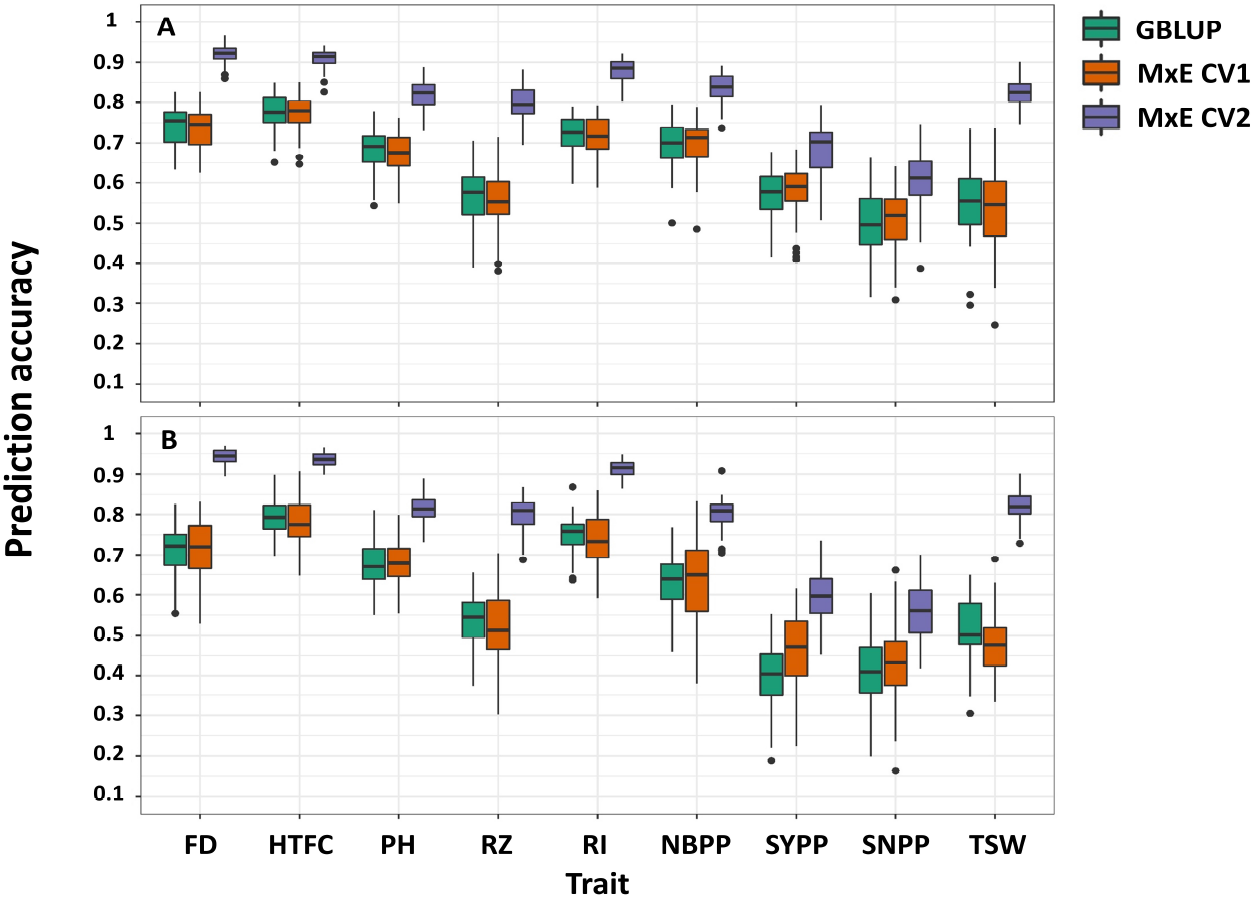
Comparison of prediction accuracies in single- and multi-environment models for predicting new lines that are not phenotyped in any environment (CV1) and predicting lines that were evaluated in only one environment (CV2) in 2018 (A) and 2020 (B) growing seasons. Flowering date (FD), height to the first capsule (HTFC), plant height (PH), reproductive zone (RZ), reproductive index (RI), number of branches per plant (NBPP), seed-yield per plant (SYPP), seeds number per plant (SNPP), and thousand-seed weight (TSW).

#### CV2 scenario

In this scenario, we evaluated the multi-environment analysis when some of the lines were not evaluated in all environments (Figure 1D). Large improvements were observed for all traits (Figure 5). The predictive accuracies of CV2 were greater than those of CV1 and the single environment GBLUP. For 2018 and 2020, improvements ranged from 17% (HTFC) to 48% (TSW) and from 15% (HTFC) to 58% (TSW), respectively. The differences in improvements were statistically significant (Table S3). Although the single-environment prediction accuracies of the yield-related traits, SYPP and SNPP, were low, using the M × E model, the gains achieved were 20% and 45% for 2018 and 20% and 28% for 2020, respectively, compared to those obtained from the single-environment analysis.

## Discussion

The future of food systems and security relies heavily on accelerating plant breeding (Lenaerts et al., 2019). Developing new varieties with high nutritional value and integration of Orphan crops such as sesame provide new opportunities to expend the human diet quality and sustainability (Dawson et al., 2019). Among the modern methods for plant breeding, genomic selection has proven effective in terms of genetic gain (Voss-Fels et al., 2019). In this study, we evaluated the genomic prediction accuracies of nine agronomic traits in sesame using a diversity panel. This was the first critical step taken toward establishing a genomic selection program for sesame.

### Performance of single-environment genomic prediction

Overall, we observed moderate-to-high prediction accuracies for all traits in the single-environment analysis (Figure 2). We did not find any significant differences between GBLUP, BayesB, BayesC, and RKHS. Variable selection methods, such as BayesB and BayesC, are expected to perform better than GBLUP in the presence of large quantitative trait locus effects (Daetwyler et al., 2010). Comparable prediction performance between GBLUP and variable selection methods supported a previous genome-wide association study reporting that only a few significant loci influenced the studied traits using the same sesame panel (Sabag et al., 2021). This suggests that agronomic traits in sesame are mostly controlled by many small-effect quantitative trait loci rather than by major quantitative trait loci. In addition, we found an association between the genomic heritability estimates and prediction accuracy. The higher the genomic heritability estimate, the higher the accuracy of genomic prediction. For example, FD and HTFC showed high genomic heritability estimates (0.72 and 0.68, respectively) and high prediction accuracies (0.72 and 0.78 on average, respectively, for both environments). Similarly, the yield components SYPP and SNPP had the lowest prediction accuracies in the two environments, as well as the lowest genomic heritability estimates. Numerous factors affect genomic prediction accuracy, such as genetic architecture, the quantitative genetic model used, trait heritability, marker density, size of the reference population, and the genetic relationship between TRN and TST (Daetwyler et al., 2010). For example, given the small sample size of the sesame diversity panel (Sabag et al., 2021), increasing the number of lines could improve the predictive performance of lowly heritable traits, such as yield components (e.g., SYPP and SNPP).

### Multi-environment analysis to enhance genomic prediction

Understanding genotype-by-environment interactions are among the main challenges for plant breeding (Cooper and DeLacy, 1994; Mathews et al., 2008). The M × E model decomposes the marker effect into the marker main effect, which borrows information from the other environment, and the marker-specific effect for each environment (Lopez-Cruz et al., 2015). No notable improvement from the M × E model was observed for CV1 when predicting the performance of new lines that were not observed in any environment. This agreed with previous reports of no strong evidence of gain in prediction for the CV1 scenario using the M × E model compared to single-environment analysis (Burguenõ et al., 2012; Lopez-Cruz et al., 2015; Crossa et al., 2016). In this scenario, no information was borrowed from the other environment. In such a case, integrating environmental covariates into the prediction model may be an alternative strategy for improving the prediction accuracy (Jarquín et al., 2014).

Many lines are often evaluated simultaneously for multiple environments in plant breeding programs (Lorenz, 2013). This leads to unbalanced field experimental designs (Lado et al., 2016), in which not all lines are present in all environments. We simulated this scenario using CV2 to investigate whether capturing environmental information improved the prediction accuracies of agronomic traits in sesame. In general, considerable improvement in prediction accuracies were observed with the M × E model compared to those of GBLUP for all traits in all environments. Our results concurred with those of previous studies (Lopez-Cruz et al., 2015; Crossa et al., 2016; Cuevas et al., 2016; Bandeira e Sousa et al., 2017; Cuevas et al., 2018), suggesting that the M × E model borrowed environmental information across environments and improved prediction accuracies (Lopez-Cruz et al., 2015). In particular, the M × E model performed well when the sample phenotypic correlations between environments were positive (Lopez-Cruz et al., 2015). This is because the covariance between any two environments is linearly related to the proportion of the genetic variance, explained by the marker main effect in the M × E model, causing the phenotypic correlation between the two environments to be positive or zero in our data. The pairs of phenotypic correlations between the environments were positive for all the agronomic traits. The mean (standard deviation) of the sample phenotypic correlation between the environments was 0.79 (0.16) (Table 1). This led to a correlation between the sample- and the ratio of variance component-based phenotypic correlations of 0.95. The positive sample phenotypic correlation between the two environments might be a critical factor in explaining why the M × E model outperformed the single-environment GBLUP model in CV2. In addition, the largest gain in prediction in CV0 compared to that in the single-environment analysis was achieved for traits with a large extent of M × E components (SNPP and SYPP) (Table 1 and Figure 4). This finding indicated that when G × E is present, the M × E model can improve prediction accuracy. Although we employed the M × E model, which only captured additive genetic effects, the extension of G × E GBLUP to RKHS has been reported to outperform G × E GBLUP in maize and wheat grain yield, especially when many environments were analyzed (Cuevas et al., 2016).

### The future of genomic prediction in a sesame breeding

Crop rotation is critical for sustainable agricultural production systems (Li et al., 2019), and the introduction of new crops, such as sesame, can be used for this purpose. Although sesame is primarily cultivated in developing countries with relatively low yields (Dossa et al., 2017), its demand for consumption is increasing. Accelerated breeding efforts are necessary to meet this growing demand. In this study, we performed genomic prediction for nine important agronomic traits in sesame using single- and multi-environment analyses for the first time. As genomic prediction is an essential first step toward the implementation of genomic selection in breeding programs, we examined the potential of using genomic prediction to enhance genetic gain in sesame while accounting for M × E. Additional improvements in yield components may be achieved using a multi-trait model along with secondary traits evaluated in this study or applying high-throughput phenotyping during the growing season (Morota et al., 2022).

## Conclusions

Currently, genetic research on sesame is limited to quantitative trait locus mapping (Teboul et al., 2020) or genome-wide association studies (Berhe et al., 2021; Sabag et al., 2021). In this study, we evaluated the usefulness of whole-genome prediction models in predicting important agronomic traits in sesame. Overall, we obtained moderate-to-high genomic prediction accuracies. Prediction performance was further enhanced by accounting for M × E. Given the reduced cost of genotyping and the availability of high-quality genomic resources for sesame, we conclude that genomic prediction has the potential to facilitate sesame breeding by transforming the prediction gain into selection decisions in Mediterranean climatic conditions.

## Author contribution statement

IS and ZP performed the field experiments. IS analyzed the data. IS drafted the manuscript. YB and GM supported IS on the data analysis. YB, ZP, and GM edited the manuscript. ZP and GM supervised the study.

## Acknowledgments

This research was supported by a Research Grant from BARD, the United States - Israel Binational Agricultural Research and Development Fund (No. IS-5400-21), the Hebrew University of Jerusalem, and Virginia Polytechnic Institute and State University. I.S. is indebted to the Samuel and Lottie Rudin scholarship foundation.

