## Supplementary material for "Multi-environment analysis enhances genomic prediction accuracy of agronomic traits in sesame": SI data

### Supplementary Materials

### Tables

| Trait | 2018 | | | 2020 | | | |
|---|---|---|---|---|---|---|---|
| | Mean | SD | CV(%) | Mean | SD | CV(%) | $H^2$ |
| Flowering date | 51.05 | 8.93 | 17.5 | 50.54 | 7.97 | 15.8 | 0.97 |
| Height to the first capsule | 71.59 | 28.66 | 40 | 92.45 | 38.74 | 41.9 | 0.96 |
| Plant height | 136.13 | 18.4 | 13.5 | 184.93 | 27.86 | 15.1 | 0.89 |
| Reproductive zone | 64.74 | 19.21 | 29.6 | 92.92 | 22.36 | 24.1 | 0.91 |
| Reproductive index | 0.48 | 0.15 | 32.7 | 0.51 | 0.14 | 28.6 | 0.97 |
| Number of branches per plant | 5.17 | 2.43 | 47.1 | 3.94 | 2.08 | 52.9 | 0.88 |
| Seed-yield per plant | 17.09 | 7.41 | 43.4 | 18.63 | 6.56 | 35.2 | 0.74 |
| Seeds number per plant | 3545 | 1558 | 44 | 6351.14 | 2204 | 34.7 | 0.66 |
| Thousand-seed weight | 3.12 | 0.5 | 16.3 | 2.94 | 0.4 | 13.6 | 0.88 |

Table S1: Mean, standard deviation (SD), coefficient of variation (CV), and heritability estimates ($H^2$) of the nine agronomic sesame traits.

| Trait | Model | 2018 | | 2020 | |
|---|---|---|---|---|---|
| | | Mean | SD | Mean | SD |
| Flowering date | GBLUP | 0.74 | 0.04 | 0.71 | 0.06 |
| | BayesB | 0.74 | 0.04 | 0.71 | 0.06 |
| | BayesC | 0.74 | 0.04 | 0.71 | 0.06 |
| | RKHS | 0.72 | 0.05 | 0.7 | 0.07 |
| Height to the first capsule | GBLUP | 0.77 | 0.04 | 0.79 | 0.04 |
| | BayesB | 0.77 | 0.04 | 0.79 | 0.04 |
| | BayesC | 0.77 | 0.04 | 0.79 | 0.04 |
| | RKHS | 0.75 | 0.06 | 0.78 | 0.04 |
| Plant height | GBLUP | 0.68 | 0.05 | 0.67 | 0.06 |
| | BayesB | 0.68 | 0.05 | 0.67 | 0.06 |
| | BayesC | 0.68 | 0.05 | 0.67 | 0.06 |
| | RKHS | 0.69 | 0.04 | 0.65 | 0.06 |
| Reproductive zone | GBLUP | 0.56 | 0.07 | 0.54 | 0.06 |
| | BayesB | 0.56 | 0.07 | 0.54 | 0.07 |
| | BayesC | 0.56 | 0.07 | 0.54 | 0.07 |
| | RKHS | 0.54 | 0.09 | 0.52 | 0.07 |
| Reproductive index | GBLUP | 0.71 | 0.05 | 0.75 | 0.05 |
| | BayesB | 0.71 | 0.05 | 0.75 | 0.05 |
| | BayesC | 0.72 | 0.05 | 0.75 | 0.05 |
| | RKHS | 0.7 | 0.07 | 0.74 | 0.05 |
| No. of branches per plant | GBLUP | 0.69 | 0.05 | 0.63 | 0.07 |
| | BayesB | 0.69 | 0.05 | 0.63 | 0.07 |
| | BayesC | 0.69 | 0.05 | 0.63 | 0.07 |
| | RKHS | 0.68 | 0.05 | 0.61 | 0.08 |
| Seed-yield per plant | GBLUP | 0.56 | 0.06 | 0.4 | 0.1 |
| | BayesB | 0.57 | 0.06 | 0.4 | 0.1 |
| | BayesC | 0.56 | 0.06 | 0.4 | 0.1 |
| | RKHS | 0.68 | 0.05 | 0.39 | 0.09 |
| Seeds number per plant | GBLUP | 0.56 | 0.06 | 0.41 | 0.1 |
| | BayesB | 0.57 | 0.06 | 0.4 | 0.09 |
| | BayesC | 0.56 | 0.06 | 0.41 | 0.1 |
| | RKHS | 0.57 | 0.07 | 0.4 | 0.08 |
| Thousand-seed weight | GBLUP | 0.55 | 0.09 | 0.51 | 0.08 |
| | BayesB | 0.55 | 0.09 | 0.51 | 0.08 |
| | BayesC | 0.55 | 0.09 | 0.51 | 0.08 |
| | RKHS | 0.55 | 0.08 | 0.5 | 0.09 |

Table S2: Single-environment genomic prediction accuracies of the best linear unbiased prediction (GBLUP), BayesB, BayesC, and reproducing kernel Hilbert spaces (RKHS) regression models obtained from repeated random sub-sampling cross-validation replicated 50 times. Mean is the mean value of the prediction accuracies, and SD is their standard deviation.

| Trait | Year | CV1 | | | CV2 | | |
|---|---|---|---|---|---|---|---|
| | | Mean | SD | P-value | Mean | SD | P-value |
| Flowering date | 2018 | 0.74 | 0.05 | 0.49 | 0.92 | 0.02 | $< .0001$ |
| | 2020 | 0.73 | 0.06 | 0.12 | 0.91 | 0.03 | $< .0001$ |
| Height to the first capsule | 2018 | 0.77 | 0.05 | 0.61 | 0.91 | 0.02 | $< .0001$ |
| | 2020 | 0.78 | 0.05 | 0.51 | 0.91 | 0.02 | $< .0001$ |
| Plant height | 2018 | 0.67 | 0.05 | 0.39 | 0.82 | 0.03 | $< .0001$ |
| | 2020 | 0.66 | 0.06 | 0.36 | 0.8 | 0.04 | $< .0001$ |
| Reproductive zone | 2018 | 0.55 | 0.08 | 0.39 | 0.8 | 0.04 | $< .0001$ |
| | 2020 | 0.52 | 0.08 | 0.23 | 0.77 | 0.05 | $< .0001$ |
| Reproductive index | 2018 | 0.71 | 0.06 | 0.47 | 0.88 | 0.03 | $< .0001$ |
| | 2020 | 0.74 | 0.05 | 0.46 | 0.89 | 0.03 | $< .0001$ |
| No. of branches per plant | 2018 | 0.7 | 0.06 | 0.81 | 0.84 | 0.03 | $< .0001$ |
| | 2020 | 0.63 | 0.07 | 0.87 | 0.78 | 0.05 | $< .0001$ |
| Seed-yield per plant | 2018 | 0.58 | 0.07 | 0.33 | 0.68 | 0.06 | $< .0001$ |
| | 2020 | 0.43 | 0.09 | 0.08 | 0.58 | 0.08 | $< .0001$ |
| Seeds number per plant | 2018 | 0.51 | 0.08 | 0.58 | 0.6 | 0.07 | $< .0001$ |
| | 2020 | 0.41 | 0.1 | 0.7 | 0.52 | 0.08 | $< .0001$ |
| Thousand-seed weight | 2018 | 0.54 | 0.1 | 0.56 | 0.82 | 0.04 | $< .0001$ |
| | 2020 | 0.48 | 0.09 | 0.07 | 0.81 | 0.03 | $< .0001$ |

Table S3: Multi-environment genomic prediction accuracies of the marker-by-environment interaction model obtained from repeated random sub-sampling cross-validation replicated 50 times. Mean is the mean value of the prediction accuracies, and SD is their standard deviation. P-value was obtained from a *t*-test comparing to the single-environment genomic best linear unbiased prediction model.
